# The PVT1, HULC, and HOTTIP expression changes due to treatment in Diffuse Large B-cell lymphoma

**DOI:** 10.1101/2024.08.05.606587

**Authors:** Milad Shahsavari, Sedigheh Arbabian, Farzaneh Hosseini, Mohamad Reza Razavi

**Author notes:** Corresponding Author: Mohamad Reza Razavi, PhD., Pasteur Institute of Iran, 69 Pasteur Avenue. 1316943551 Tehran, Iran.

## Abstract

Diffuse large B-cell lymphoma is the most common histological subtype of non-Hodgkin’s lymphomas. It is an aggressive malignancy that displays great heterogeneity in morphology, genetics, biological behavior and treatment response owing to chromatin remodeling and epigenetics.

Bioinformatic-based approaches were used to understand the possible signaling pathways of the three lncRNAs *PVT1*, *HULC*, and *HOTTIP*. Furthermore, their expression levels were quantitatively evaluated in 100 patients before and after the treatment.

The results revealed that gene expression was significantly upregulated in *PVT1*, *HULC*, and *HOTTIP* by 7.39±8.48-, 5.924±7.536-, and 4.137±5.863 fold, respectively, relative to normal cases. Post-treatment measurement of lncRNA expression indicated that *PVT1* and *HOTTIP* were significantly downregulated.

Interestingly, the expression levels of *PVT1*, *HULC*, and *HOTTIP* were significantly higher in DLBCL patients aged > 60 years than in those aged < 60 years. In addition, there was a significant positive correlation between *HULC* and *HOTTIP* expression.

The analysis of overexpressed lncRNA-miRNA interaction indicated different deregulated miRNA targets and the protein targets of upregulated lncRNAs are mainly with histone modification, DNA methylation/demethylation, and protein methyltransferase activity.

**Summary blurb:** The lncRNAs *PVT1*, *HULC*, and *HOTTIP* expression is significantly upregulated before treatment and reduce to normal level after treatment. It can be used as diagnostic marker or prognostic means especially in Relapsed/refractory DLBCL.

## Introduction

Diffuse large B cell lymphomas (DLBCLs), neoplasms of medium or large B lymphoid cells, account for 25 to 35 percent of all non-Hodgkin lymphomas. The standard therapy for DLBCL patients is a phase III trial with a standard regimen including rituximab, cyclophosphamide, doxorubicin, vincristine, and prednisone known as R-CHOP (Coiffier et al., 2010).

In most cases, R-CHOP has been shown to enhance the 5-year overall survival of patients. However, a significant proportion of patients, approximately 30 to 50 percent, do not achieve a cure with this regimen, which compromises patients suffering from primary refractory disease and patients with DLBC relapse after attaining complete remission (Coiffier & Sarkozy, 2016). Recently, novel therapy has introduced Lenalidomide plus R-CHOP (R2CHOP) as a promising immunomodulatory combination to enhance outcomes in newly diagnosed DLBCL patients (Desai et al., 2020). Therefore, a comprehensive understanding of DLBCL biology and behavior is crucial for exploring alternative treatment approaches.

Recent studies have increasingly focused on exploring the molecular underpinnings and mechanisms of DLBCL to facilitate more effective management (Danilov et al., 2022). Three distinct molecular subtypes of DLBCL were identified, namely germinal-center B-cell-like (GCB) DLBCL, activated B-cell-like (ABC) DLBCL, and primary mediastinal B-cell lymphoma (PMBL), using high-throughput sequencing (Roschewski et al., 2020). Notably, the most frequent genetic alterations in DLBCL are associated with chromatin and epigenetics (Bakhshi & Georgel, 2020; Harrop et al., 2022), and researches indicate that long non-coding RNAs (lncRNAs) are frequently dysregulated in cancer, including DLBCL (Pathania, 2023).

LncRNAs are transcripts with lengths over 200 nucleotides; they mostly remain in the nucleus after transcription. Numerous functions have been proposed for lncRNAs through interactions with DNA, RNA, or proteins (Cao, 2014). LncRNAs act as tumor suppressors or oncogenes and participate in diverse signaling pathways (Do & Kim, 2018). Noteworthy, the study of lncRNA expression in different types of cancer could be utilized as promising biomarkers and probabilistic therapeutic targets (Beylerli et al., 2022; Mohammadian et al., 2024). Hence, a precise understanding of DLBCL biology and its function in transcriptional regulation by chromatin modifications could be crucial in managing this malignant and fatal disease.

Building on previous literature, three crucial lncRNAs, PVT1 (Plasmacytoma Variant Translocation 1) (Yang et al., 2020), HULC (Highly Up-regulated in Liver Cancer) (W. Peng et al., 2016), and HOTTIP (HOXA transcript at the distal tip) (Habieb et al., 2022), were quantitatively investigated and evaluated in one hundred Iranian patients before and after conventional treatments, which is novel compared to the previous studies. Subsequently, advanced bioinformatic-based approaches were employed to comprehend the potential signaling pathways and biological function of these lncRNAs.

## Results

### Demographic and Clinicopathologic information of patients

The mean age±SD of the studied patients was 63.96±12.34 years; 53% were male, and 47% were female. Also, the mean age±SD of the control group was 61.47±12.03; 50% were male, and 49% were female.

Twelve patients discontinued treatment during the follow-up. All eligible patients in this study were screened for other types of malignancy (Table 1).

**Table 1.** The demographic and clinicopathological information of DLBCL patients. CR stands for complete response; Cru stands for complete response (Uncomfrimed); PD stands for progressive disease; PR stands for partial Response.

| Factors | DLBCL Patients (n=100) | Control Group (N=55) |
| --- | --- | --- |
| Age Mean±SD (year) | 63.96±12.34 | 61.47±12.03 |
| Age range (year) | 45-84 | 41-82 |
| Male | 52 | 28 |
| Female | 48 | 27 |
| Stage 1 lymphoma | 17 | NA |
| Stage 2 lymphoma | 20 | NA |
| Stage 3 lymphoma | 38 | NA |
| Stage 4 lymphoma | 25 | NA |
| With symptoms of lymphoma | 54 | NA |
| Without symptoms of lymphoma | 46 | NA |
| CR | 39 | NA |
| Cru | 18 | NA |
| PD | 15 | NA |
| PR | 28 | NA |
| Relapse | 64 | NA |
| No relapse | 24 | NA |
| Ceased | 12 | NA |

### The expression of lncRNAs during the treatment process

The expression of lncRNAs was checked on 100 snap-frozen tissue before the conventional treatment of DLBCL. Interestingly, the mean±SD of PVT1 relative expression compared to the normal cohort was 7.412 ± 2.497, significantly (p <0.001) higher than the normal group. The mean±SD of PVT1 relative expression of PVT1 after treatment (2.94 ± 1.22) was significantly (p <0.001) decreased compared to the relative expression of PVT1 before treatment. Noteworthy, there was no significant difference (p =0.053) in the relative expression of PVT1 after treatment compared to the normal cohort. (Figure 1-A)

**Figure 1.**
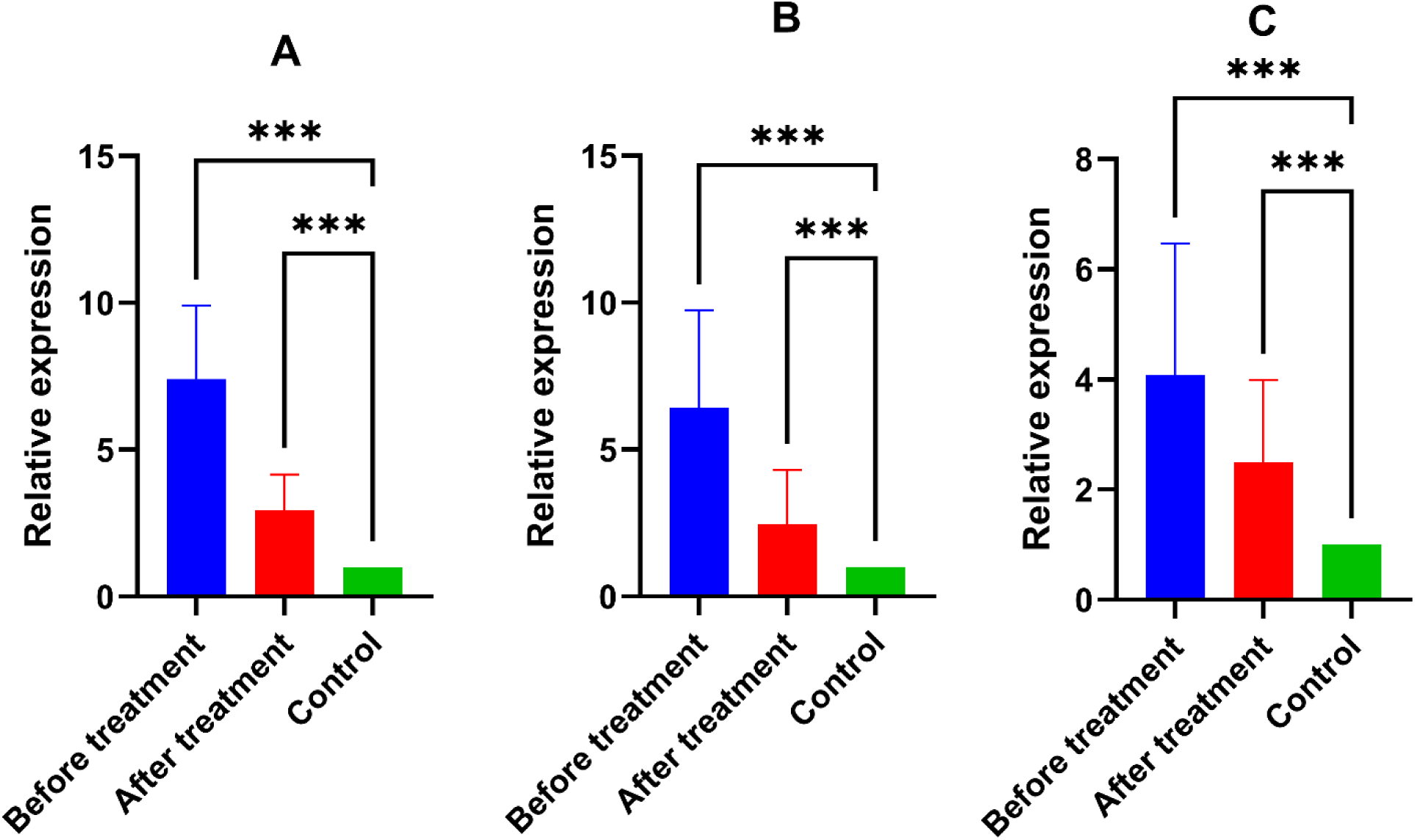
The relative expression of lncRNAs. A) The expression of PVT1 in DLBCL patients before treatment is significantly (p <0.001) higher compared to the normal control also, after treatment, it significantly (p <0.001) decreased, while the expression of PVT1 after treatment is not altered statistically significantly compared to the normal cohort. B) The expression of HULC in DLBCL patients before treatment is significantly (p <0.001) higher compared to the normal control, and also, after treatment, significantly (p <0.001) decreased. In contrast, the expression of HULC after treatment is not altered statistically significantly compared to the normal cohort. C) The expression of HOTTIP in DLBCL patients before treatment is significantly (p <0.001) higher than in the normal control, while the expression after treatment compared with before treatment decreased less than other lncRNAs (p <0.032). *** means p <0.001; * between the HOTTIP expression of control and after treatment means p < 0.31; * between the HOTTIP expression of before treatment vs. after treatment means p <0.32; ns: insignificant.

Furthermore, the mean±SD of HULC relative expression was 6.42 ± 3.32, significantly (p <0.001) higher than the normal group. Likewise, the relative expression of HULC in patients after treatment (2.46 ± 1.85) was significantly (p =0.053) decreased compared to the HULC relative expression before treatment, and there was no significant difference (p =0.162) compared to the normal group. (Figure 1-B)

Additionally, the mean±SD of HOTTIP relative expression (4.09 ± 2.38) was significantly higher in DLBCL patients (p < 0.001). After treatment, it was significantly downregulated (2.50 ± 1.49) (p =0.032). In contrast with the other studied lncRNAs, there was a significant (p =0.031) difference between the relative expression of HOTTIP in DLBCL patients after treatment compared to the normal cohort. (Figure 1-C)

Noteworthy, the considerable differences in the SD mentioned above are due to the outlier expression results in the data. Hence, we investigated the correlation of the expression level with various factors. Interestingly, this study found that PVT1, HULC, and HOTTIP expression was significantly (p <0.001) higher in DLBCL patients above 60 than in patients below 60. (**Figure 2-A, B, C**) The disease-free survival analysis found no significant difference (p =0.3804) between the patients above 60 and patients below 60. (**Figure 2-D**)

**Figure 2.**
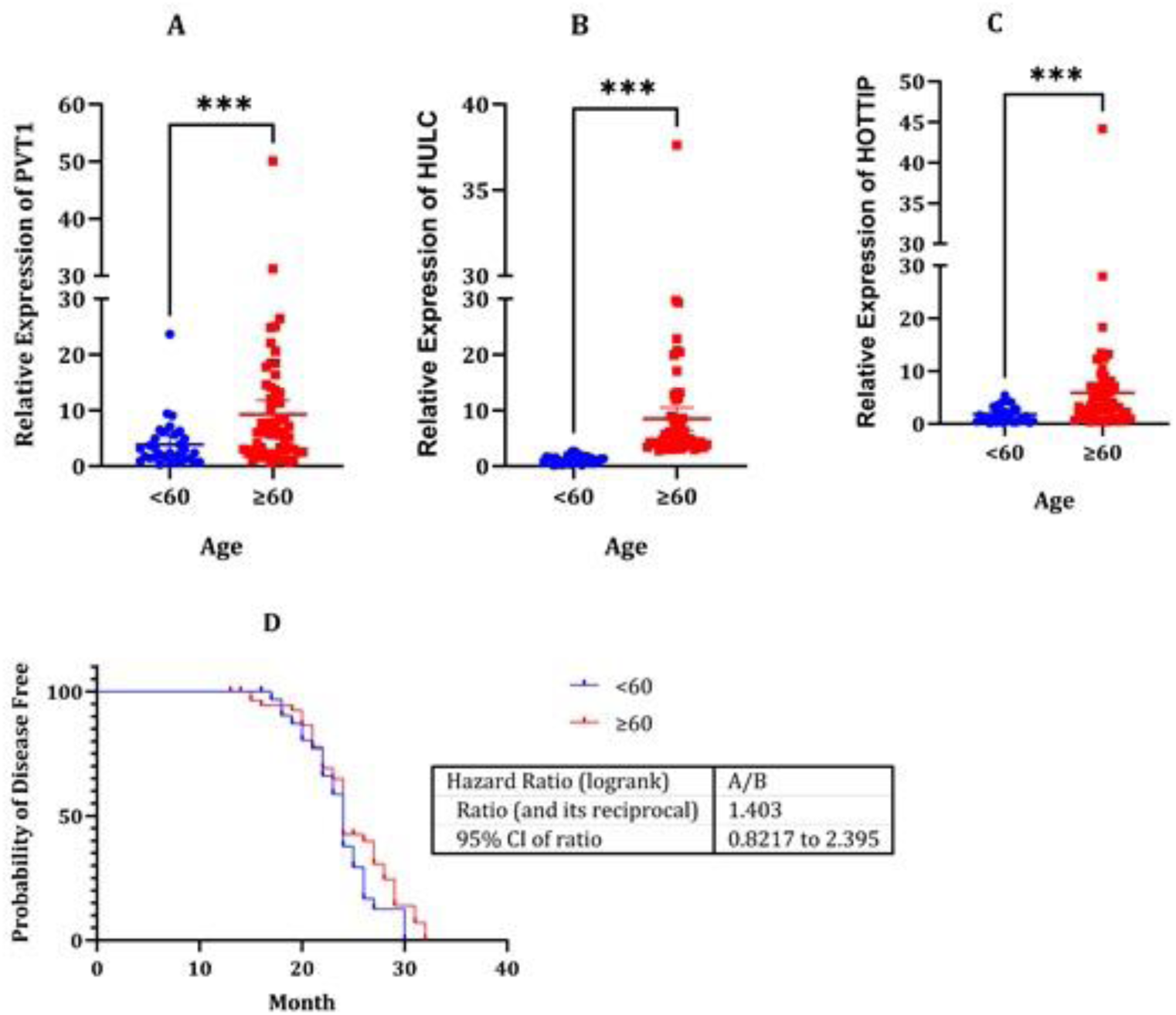
The relative expression of lncRNAs between the DLBCL patients (before treatment) with age above 60 compared to patients below 60. The expression of A) PVT1, B) HULC, and C) HOTTIP in DLBCL patients between two age groups demonstrates that these three lncRNAs have a significant (p< 0.001) higher expression in the DLBCL patients over 60 years old. D) Disease-free survival analysis reveals that these expressions do not impact patients’ disease-free state based on age (p =0.3804).

The study’s results indicate that the expression pattern of the studied lncRNAs gradually increased with age (Figure 3). The expression pattern of PVT1 gradually increased, while this increase was insignificant (p= 0.084) among age groups. Furthermore, the expression pattern of HULC and HOTTIP was significantly (p< 0.001) increased among the age groups. Noteworthy, it was found a significant positive correlation (Pearson r= 0.3669, p< 0.001) between the expression HULC and HOTTIP (**Figure 3**).

**Figure 3.**
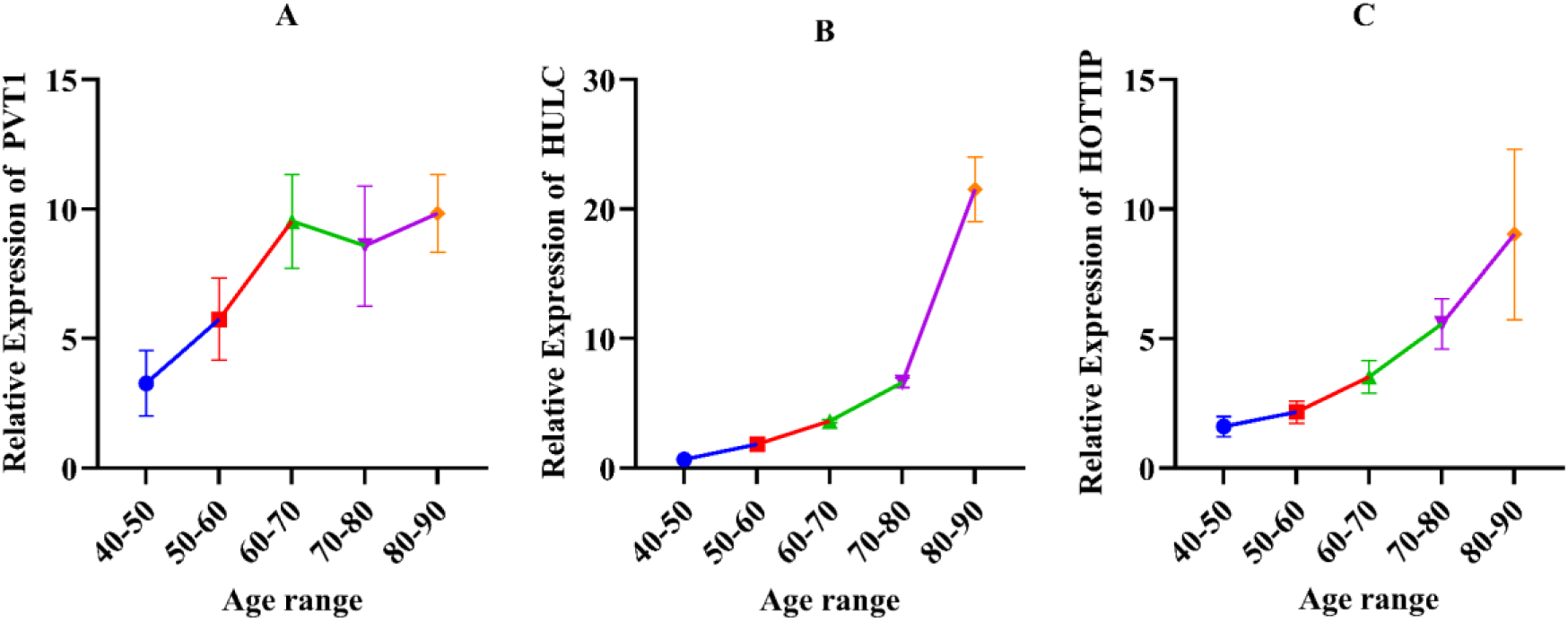
The relative expression of studied lncRNAs PVT1, HULC, and HOTTIP in the age groups. A) PVT1, B) HULC, and C) HOTTIP relative expressions demonstrate an ascending trend in the expression of PVT1, HULC, and HOTTIP in DLBCL patients related to their age.

### The cut-off for expression levels of PVT1, HULC, and HOTTIP after treatment

ROC analysis was used to estimate the predictive accuracy of the expression level lncRNAs after the duration of treatment. The findings demonstrated that the expression levels of PVT1 with cut-off < 4.210 with Sensitivity 87.50% (95% CI: 78.99% to 92.87%), Specificity 90.00 (95% CI: 82.56% to 94.48%), HULC with cut-off < 4.305 with Sensitivity 84.09% (95% CI: 75.05% to 90.28%), Specificity 72.00 (95% CI: 62.51% to 79.86%), and HOTTIP with cut-off < 2.680 with Sensitivity 64.77% (95% CI: 54.37% to 73.94%), Specificity 73.00 (95% CI: 63.57% to 80.73%), could reflect the effective treatment.

**Figure 4.**
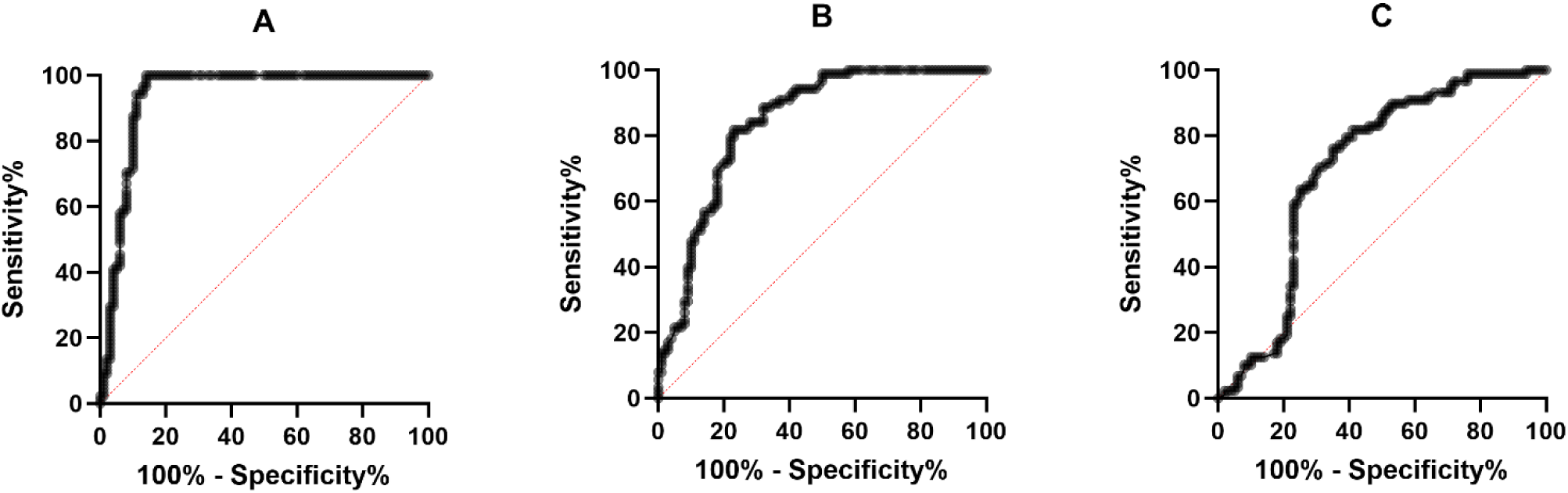
ROC curve for PVT1, HULC, and HOTTIP. A) PVT1, B) HULC, and C) HOTTIP. ROC, Receiver-operating characteristics

### Bioinformatic analysis

#### Gene ontology

Combining putative interactors with Gene ontology analysis shows that the putative targets of these lncRNAs have critical functions in regulating mRNA processes, including mRNA mediating catabolic and metabolic processes and negative regulation of gene expression (**Figure 5**). The miRNAs deregulate the translation of mRNA transcripts of protein-coding genes. The repression can occur by binding to the transcript or promoting its degradation. It has been proposed that miRNAs can regulate the expression of protein-coding genes at the post-transcriptional level (Madhumita & Paul, 2022). Hence, by GO enrichment analysis, the putative signaling networks of regulated mRNAs could be revealed (Hueso et al., 2020) (**Figgure S**1).

The first step of identifying the putative signaling of PVT1, HULC, and HOTTIP was constructing RNA-protein interaction. All putative targets were obtained from the RNA interactome repository. Interestingly, the network demonstrates that all three lncRNA directly interact with MOV10. MOV10 is an RNA helicase associated with the RNA-induced silencing complex component Argonaute (AGO), likely resolving RNA secondary structures (Nawaz et al., 2022). Also, the studied lncRNAs directly interact with EZH2, LATS2, AGO3, AGO4, FBL, STAT1, and AGO2. (Figure 5).

**Figure 5.**
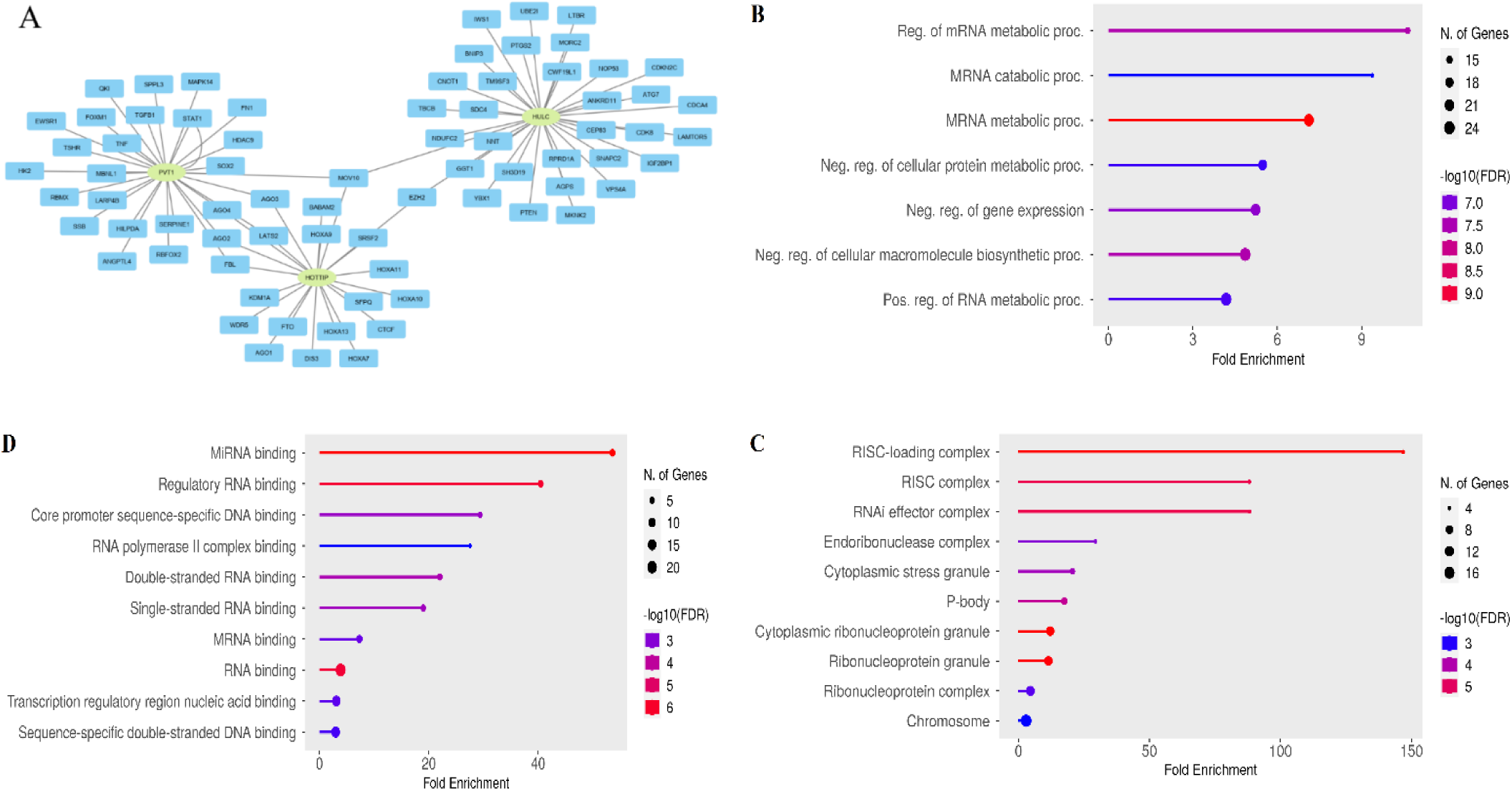
Interaction network, Gene Ontology of the potential targets. All potential targets were combined for the interaction network. Long non-coding RNAs (lncRNAs) are represented by green ellipses, while RNA-binding proteins are depicted as blue rectangles. Figures B to D each display the top ten pathways from the Gene Ontology enrichment analysis of potential targets. The Y-axis denotes the biological process, cellular component, and molecular function, while the X-axis indicates the enrichment score. In the KEGG analysis, the pathway name is on the Y-axis, and the enrichment score is on the X-axis. The bubble size signifies the number of genes involved in the pathway enrichment. The color, which ranges from red (smaller p-value) to blue, is based on the p-value (false discovery rate (FDR)). Figure A shows the interaction network among PVT1, HULC, and HOTTIP and their potential RNA-binding proteins. Figures B, C, and D represent the biological process, cellular component, and molecular function.

#### LncRNA-miRNA-mRNA interaction

The lncRNA-miRNA interactions were analyzed using the RNAInter database (http://rnainter.org/). The studied lncRNAs, PVT1, HULC, and HOTTIP, were predicted to interact with 210 miRNAs. By analyzing the lncRNA-miRNA interaction network with the Cytoscape, it was demonstrated that three lncRNAs have interactions with hsa-miR-30b-5p, hsa-miR-203a-3p, and hsa-miR-15a-5p. The interacting miRNAs with HOTTIP, with a confidence score of more than 0.5, are hsa-miR-30b-5p and hsa-miR-216a-5p. The HULC interacting miRNAs with a confidence score of more than 0.5 are hsa-miR-200a-3p and hsa-miR-372-3p. The interacting miRNAs with PVT1 with confidence score above 0.5 are hsa-miR-186-5p, hsa-miR-195-5p, hsa-miR-200b-3p, hsa-miR-26b-5p, hsa-miR-200c-3p, hsa-miR-152-3p, hsa-miR-488-3p, hsa-miR-200a-3p, hsa-miR-1207-5p and hsa-miR-203a-3p. For each lncRNA, the interacting miRNAs with higher confidence scores are shown in **Supplement 3 and Figure 6.**

**Figure 6.**
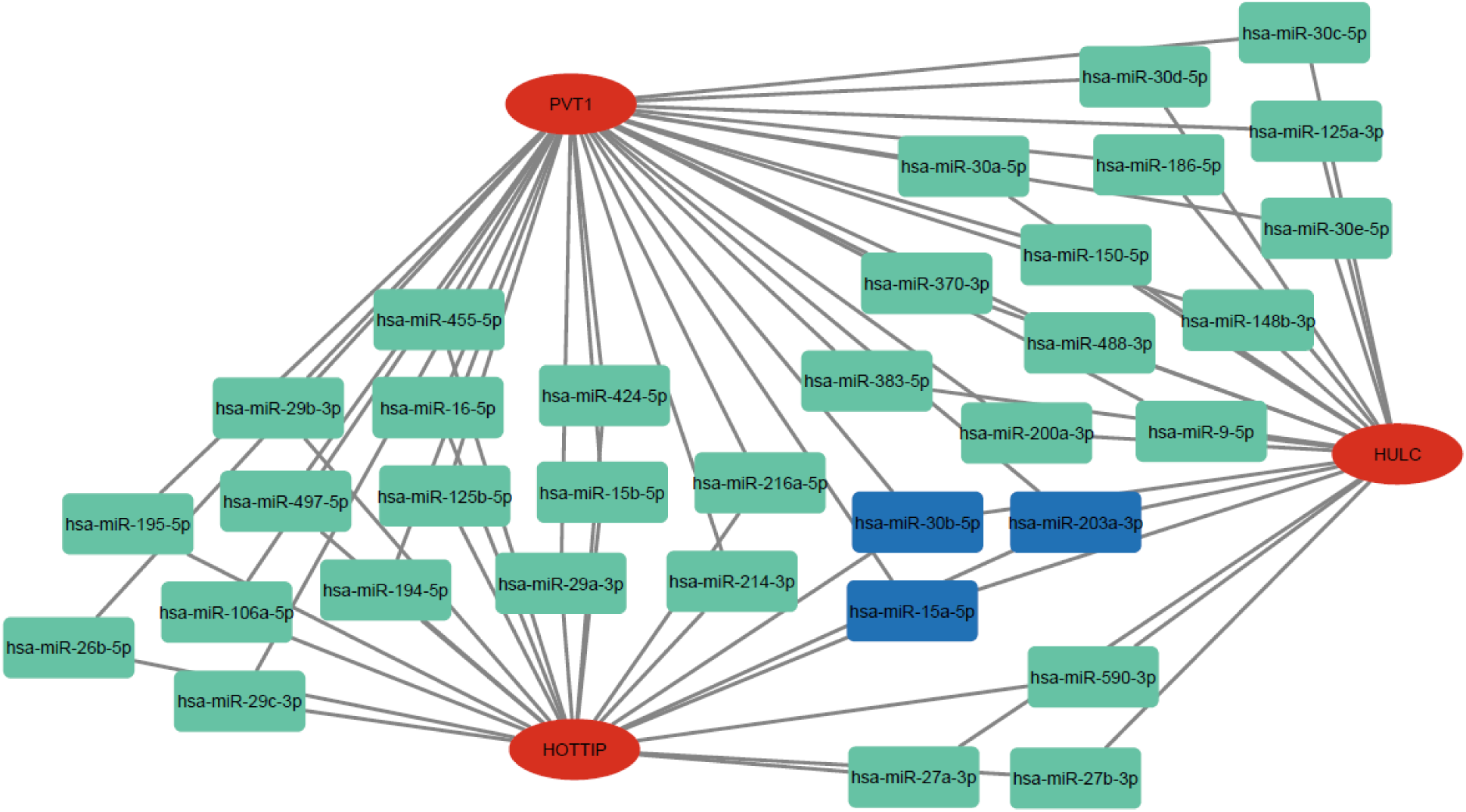
LncRNA-miRNA Interaction Network. The figure presents the interaction network between long non-coding RNAs (lncRNAs) and microRNAs (miRNAs), as analyzed using Cytoscape. The focus is on three specific lncRNAs: PVT1, HULC, and HOTTIP, which are predicted to interact with 210 miRNAs. Notably, these lncRNAs have interactions with hsa-miR-30b-5p, hsa-miR-203a-3p, and hsa-miR-15a-5p. The figure demonstrates the role of lncRNAs as miRNA sponges, binding to miRNAs and thereby reducing their regulatory effect on messenger RNAs (mRNAs).

## Discussion

This study investigated the expression of PVT1, HULC, and HOTTIP in DLBCL in one hundred patients before and after conventional treatment. Also, the expression level was compared with the normal control group. Unfortunately, eighteen patients ceased during the two-year follow-up. All three lncRNAs were upregulated (p <0.001) in newly diagnosed patients, which could be deciphered as the possible role of oncogenesis. After the treatment, the expression of PVT1 and HULC was downregulated toward the mean expression of the control group, and there was no significant difference between the DLBCL patients and the control group. They are following the results of Yang et al., who demonstrated that the expression of PVT1 is upregulated in DLBCL patients and positively associated with adverse clinicopathological outcomes (Yang et al., 2020). Furthermore, Peng et al. claimed that HULC is upregulated in DLBCL patients, and HULC knockdown could induce growth arrest and cell apoptosis in vitro (Wei Peng et al., 2016).

Additionally, the expression of HOTTIP was downregulated in DLBCL patients after treatment (p = 0.032) related to the newly diagnosed patients. Interestingly, Habeib et al. conveyed that the serum level of HOTTIP is elevated in DLBCL patients (Habieb et al., 2022). Another intriguing study result was the over-expression of PVT1, HULC, and HOTTIP (p <0.001) in patients over 60 years old compared to patients below 60 years old. In contrast, the expression of PVT1, HULC, and HOTTIP in patients below 60 years old was significantly (p <0.001) higher than in the control group. Studies suggest that lncRNA expression may contribute to or reflect the tissue-specific fine-tuning of the aging-associated process. (Marttila et al., 2020) and cause different expression levels of three lncRNAs in DLBCL patients depending on their age.

The ROC analysis confirms that the expression levels of PVT1 (< 4.210) have a sensitivity of 87.50% and a specificity of 90.00. Similarly, HULC (< 4.305) shows a sensitivity of 84.09% and specificity of 72.00, while HOTTIP (< 2.680) indicates a sensitivity of 64.77% and specificity of 73.00. These findings suggest that these lncRNAs could reflect the effectiveness of the treatment. These findings indicate that the expression levels of these lncRNAs may serve as valuable biomarkers for predicting treatment outcomes in patients.

Afterward, the miRNA-mRNA interaction network showed that hsa-miR-30b-5p, hsa-miR-203a-3p, and hsa-miR-15a-5p could interact with 1441 mRNAs and regulate their expression levels. GO enrichment analysis revealed the putative signaling networks of regulated mRNAs. The BP enrichment showed that most putative mRNA targets regulate RNA metabolic processes, macromolecule biosynthetic processes, and transcription.

In a recent study, the expression of *miR-195* decreased, and the expression of *miR-125* increased in patients with DLBCL (Li et al., 2023). The results of the current study reveal the deregulation of hsa-miR-125b-5p by HOTTIP, hsa-miR-125a-3p by HULC, and hsa-miR-195-5p, hsa-miR-125b-5p and hsa-miR-125a-5p by PVT1. The role of miR-195 sponging in tumorigenesis and immune escape has been shown in large B-cell lymphoma (Chen et al., 2020; Wang et al., 2019). It is indicated that MALAT1, PD-L1, and CD8 were upregulated while miR-195 was down-regulated in DLBCL tissues (Wang et al., 2019). One of the detected targets of HULC and PVT1 is mir*-203,* and it has been revealed that MIR-203 methylation is tumor-tissue specific in DLBCL (Voropaeva et al., 2022).

By using miRNA target prediction, it has been revealed that the miR-30 family was involved in Ibrutinib resistance in ABC-DLBCL and that the miR-30 family can directly down-regulate BCL6 (Li et al., 2021). Recent scientific evidence has revealed that long non-coding RNAs (lncRNAs) are epigenetic mediators in cancer initiation and development. The interaction between lncRNA and miRNA is a well-known phenomenon, and it may lead to the development of malignancy and resistance to cancer treatment. Also, the lncRNA-miRNA axis plays a crucial role in regulating cell death by apoptosis and autophagy in various types of cancer. In addition, it can impact other cancer signaling pathways like PI3K/Akt, STAT3, Wnt/β-catenin, and EZH2 (the main H3K27 KMT) (Statello et al., 2021).

The expression clustering analysis has revealed two distinct epigenetic-related clusters, EC1 and EC2. The EC1 cluster comprises TP53, MYD88, HIST1H1D, HIST1H1C, KMT2D, and EZH2 mutations, which regulate DNA methylation/demethylation, histone methyl transferase activity, and protein methyl transferase activity. The second cluster accompanied B2M, CD70, and MEF2B mutations that are associated with DNA damage repair, immune signaling, increased CD8+ T-, γδT-and T helper cells, and immunogenic cell death modulators (Wang et al., 2022). The results reveal that TP53 and EZH2 are molecular targets of elevated lncRNAs in DLBCL patients.

The molecular targets, MYC, BCL2, TP53, CDKN1A, and CDKN2B, are proteins relevant to GC B-cell and DLBCL physiology, and EZH2 is relevant to GC B-cell and DLBCL epigenetics (Bakhshi & Georgel, 2020)According to our in silico analyses, H3K4 and H3K27 methyl transferases are the main targets of three studied lncRNAs, including PVT1, HULC, and HOTTIP. The targets’ Molecular function is mainly to modify Histone H3, instruct transcriptional regulation, and represent an epigenetic code.

Our findings indicate that PVT1, HULC, and HOTTIP could interact with RNA-binding proteins such as EZH2, MOV10, LATS2, AGO1, FBL, SUZ12, STAT1and STAT4. These proteins are involved in regulating mRNA metabolic processes, and they modulate the transcription and translation of mRNA (Dreyfuss et al., 2002). Also, the study indicates Lysine acetyltransferases type2 and CAT2 deregulated by HOTTIP. It was reported that Histone acetyltransferase KAT2A is closely related to histone acetylation regulators, and it was overexpressed in DLBCL patients with poor prognoses (Yu et al., 2023).

Polycomb Repressive Complex 2 (PRC2) plays a crucial role in mammalian development through its temporospatial repression of gene expression. The catalytic subunit of PRC2, EZH1, or EZH2 methylates histone H3 lysine 27 in forms of mono-, di-, and tri-methylation (H3K27me1/2/3). H3K27me2/me3 characterizes facultative heterochromatin. PRC2 has a pleiotropic role in cancer. Most cancers are associated with gain-of-function mutations or the overexpression of EZH2, while PRC2 loss-of-function mutations in EZH2, EED, or SUZ12 can also be linked with cancer. In general, aberrant PRC2 activity changes global histone H3K27 methylation, reducing facultative heterochromatin and gaining euchromatin. This leads to the activation of oncogenes and/or the inactivation of tumor suppressor genes, which stimulates carcinogenesis (35). The gene expression changes mediated by EZH2/miRNA interactions enable cancer cells to out-compete their neighboring normal tissues and are passed on from one cancer cell to the next.

During oncogenesis, EZH2 interacts with other miRNAs and critically dysregulates multiple cyclin-dependent kinases (CDKs), including p16INK4A (CDKN2A), p14ARF (CDKN2A) and p21Cip1 (CDKN1A) (Hillyar et al., 2022). Investigations have revealed the importance of EZH2 in maintaining physiological balance. EZH2 encodes a histone methyltransferase that causes transcriptional repression and acts as an oncogene, promoting the development and progression of various human malignancies, including DLBCL (Chung, 2022). It plays a vital role in developing the lymphoid system and its deregulation due to genetic or non-genetic causes, can result B cell- and T cell-related lymphoma or leukemia (Li & Chng, 2019). LncRNAs act as a scaffold for EZH2 and are noticeable regarding EZH2-mediated oncogenesis (Tsai et al., 2010). Hence, it could be deciphered by targeting the lncRNAs, the pathogenesis of EZH2 could be modulated, and promising therapeutic agents may be developed.

Interestingly, The PVT1, HULC, and HOTTIP directly and indirectly regulate the Wnt/β-catenin pathway, which is critical in embryonic development and adult tissue homeostasis. The deregulation of Wnt/β-catenin signaling often leads to serious diseases, and PVT1 promotes DLBCL progression (Tao et al., 2022).

The current study revealed that deregulate FOXA1 and FOXA2, FOXP1 and FOXM1 by HOTTIP, FOXA1, FOXM1and FOXA2 by PVT1, and FOXA2 and FOXA1 by HULC. It has been shown that increased abundance of FOXP1 in DLBCL is a predictor of poor prognosis and resistance to therapy (Walker et al., 2015). Besides, HULC could promote the proliferation and invasion of pancreatic cancer cells and promote apoptosis via the Wnt/β-catenin signaling pathway (Ou et al., 2019). Aligned with the recent investigations on the putative role of PVT1 and HULC, studies demonstrate that HOTTIP plays an essential role in tumor response to chemotherapy via activating the Wnt/β-catenin pathway (Li et al., 2016).

Also, it has been shown that the expression of the PVT1 is suppressed by tumor suppressor protein p53, leading to the inactivation of the TGF-β/Smad signaling pathway and exerting an anti-oncogenic function on glioma development (Takahashi et al., 2021).

In conclusion, the primary function of lncRNAs is chromatin regulation. The instability of nuclear lncRNAs indicates their involvement in regulating gene expression, fine harmonizing in response to stimuli, and participation in the turnover of transcription factors. The study’s findings indicate that PVT1, HULC, and HOTTIP lncRNAs have diverse roles in cellular biology, cancer progression, and treatment response, and intend to understand the exact mechanism of these lncRNAs-miRNA-mRNA interactions in carcinogenesis and treatment response. The conditional specific expression patterns of PVT1, HULC, and HOTTIP lncRNAs suggest they can be used as diagnostic biomarkers or prognostic means of treatment response in different age groups. The results emphasize that potentially using the wide-spectrum inhibitors of histone lysine methyltransferases or acetyltransferases would be a promising approach for targeting epigenetic and chromatin biology deregulation in diffuse large B-cell lymphoma.

## Materials and Methods

### Tissue Samples Collection

The study involved expressing lncRNAs on 240 snap-frozen lymph tissue specimens. The specimens included 100 DLBCL tissues before treatment, 88 lymph tissues of DLBCL patients after complete treatment, and 58 normal tissues as controls. Twelve patients were deceased during the study.

The pathologists and physicians meticulously collected patients’ demographic and clinicopathological characteristics for further analysis. All patients provided written consent. The study’s eligibility criteria for the DLBCL patients were based on the clinician and pathologist’s diagnosis using the Immunohistochemistry method (IHC) and physician diagnosis. The control cohort, on the other hand, had no types of malignancy or inflammation. This was confirmed by evaluating conventional tumor markers, including Carcinoembryonic Antigen (CEA), Carbohydrate Antigen 19-9 (CA 19-9), Cancer Antigen 125 (CA 125), Cancer antigen 15-3 (CA 15-3), total Prostate-Specific Antigen (PSA), Beta-Human Chorionic Gonadotropin (BHCG), and Alpha Feto Protein (AFP). It should be noted that CA 125 was evaluated in female patients, and total PSA was assessed only in male patients. All evaluations were based on the electrogenerated chemiluminescence (ECL) method. (**Table S1**).

### RNA Extraction and Relative Quantitative Real-time PCR Analysis

Whole RNAs were extracted from tissues using the RNA extraction kit. The extracted RNA’s quantity and quality were evaluated by spectrophotometer (Thermo Scientific™ NanoDrop™) with the 260/280 ratio of ∼2.0 and agarose 1% gel electrophoresis.

All extracted RNA from samples was synchronized to 1000 ng/mL and reversely transcribed into cDNA using the QuantiTect Reverse Transcription kit (QIAGEN). qRT-PCR was performed using Master Mix 2x (®Primer Design) in the Eco Illumina machine (Illumina, Inc.). Primer sequences are listed in Table 2.

**Table 2.**
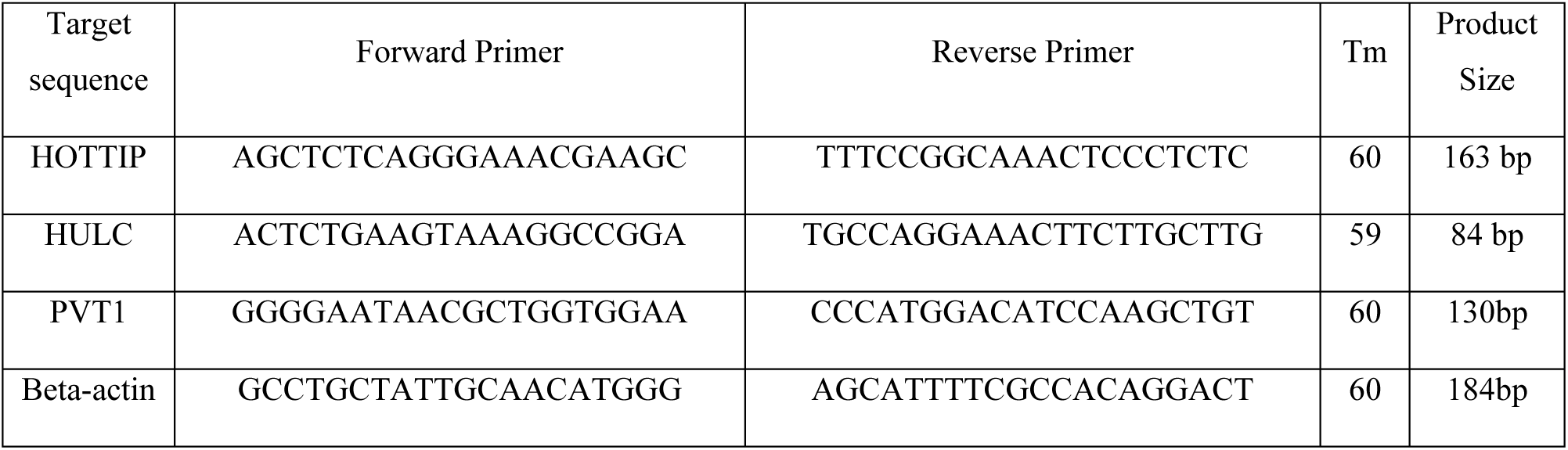
The Primer nucleotide sequences are used to validate gene expression.

The expression of studied lncRNAs was evaluated in three independent repeats, and the mean value was used for comparative analyses. The relative expression of studied lncRNAs is calculated against the expression level of Beta-actin using the 2^−ΔΔCt^ method (13).

### *In Silico* Data Analysis

#### Gene Ontology Enrichment Analysis

The gene ontology (GO) enrichment analysis was performed with the ShinyGO 0.77 database (http://bioinformatics.sdstate.edu/go/) to identify the functions of identified modules (with an FDR cut-off < 0.05). The biological properties of the selected genes were characterized through three sub-ontologies: biological process (BP), cellular component (CC), and molecular function (MF).

#### Construction of lncRNA-RNA-binding proteins interaction network

This study investigated the putative signaling pathways of PVT1, HULC, and HOTTIP by defining the potential targets of these three lncRNAs. For this purpose, we used the RNAInter database (14). The interaction type was determined to be RNA-protein interactions, with a confidence score ranging from 0.1 to 1.0.

#### Construction of lncRNA-miRNA and miRNA-mRNA interaction network

The prediction of lncRNA-miRNA interactions was performed using the RNAInter database (http://rnainter.org/). Common miRNAs were identified by analyzing the lncRNA-miRNA network through the Cytoscape network analyzer. miRNAs with a higher degree of interaction (three) were selected to identify the miRNA-mRNA interaction. Afterward, the Mirdb (http://mirdb.org/) was utilized to determine the putative mRNA targets of selected miRNA.

### Statistical Analysis and Validation

The statistical analyses were conducted using GraphPad Prism 9.3.1. The Mann-Whitney U test was used to determine differences between groups, while the One-way Analysis of Variance (ANOVA) with Benjamini-Hochberg correction identified differences among the three groups. The threshold probability for statistical significance was less than 0.05. Pearson’s correlation coefficient was used to determine the association between the expression of lncRNAs and measured factors. A Receiver Operating Characteristic (ROC) curve was generated to determine optimal sensitivity, specificity, and cut-off levels of lncRNAs, enabling more accurate and reliable predictions using lncRNAs as biomarkers.

## Acknowledgments

We gratefully acknowledge Miss. Mariam Amin is responsible for supporting medical writing and editing.

## Authors contributions

Milad Shahsavari: Wrote the first draft of the manuscript and contributed to the conception, design, experiments, and data analysis. Also, performed bioinformatics analysis, sampling, and laboratory experiments.

Sedigheh Arbabian: Contributed to the conception, design, experiments, and data analysis. Farzaneh Hosseini: Contributed to the conception, design, experiments, and data analysis. Mohamad Reza Razavi: Contributed to the conception, design, experiments, and data analysis and rewrote and revised the final manuscript. Also, performed bioinformatics analysis, sampling, and laboratory experiments.

## Ethics approval and consent to participate

The study conducted at Tehran Islamic Azad University, Tehran North Branch, was approved by the Ethics Committee, with approval ID: IR. IUA.TNB. REC: 1401. 040. The research team prioritized the safety and well-being of all participants, meticulously following all necessary guidelines and regulations.

Before participating, all participants were informed about the study’s purpose and procedures and provided with an informed consent form. The form clearly outlined that their clinical samples and personal data would be used for the study under their physician’s supervision. The research team took extra care in preparing the informed consent form; all participants or legal guardians willingly signed it before proceeding. Participants were free to withdraw their consent at any point during the study. Throughout the study, the research team upheld the highest ethical standards and strictly followed all relevant regulations to guarantee the privacy and confidentiality of all participants’ data.

